# Membrane free-energy landscapes derived from atomistic dynamics explain nonuniversal cholesterol-induced stiffening

**DOI:** 10.1101/2023.02.02.525347

**Authors:** Giacomo Fiorin, Lucy R. Forrest, José D. Faraldo-Gómez

**Affiliations:** National Institute for Neurological Disorders and Stroke, Bethesda, MD, USA; National Heart, Lung and Blood Institute, Bethesda, MD, USA

## Abstract

All lipid membranes have inherent morphological preferences and resist deformation. Yet adaptations in membrane shape can and do occur at multiple length scales. While this plasticity is crucial for cellular physiology, the factors controlling the morphological energetics of lipid bilayers and the dominant mechanisms of membrane remodeling remain unclear. An ongoing debate regarding the universality of the stiffening effect of cholesterol underscores the challenges facing this field, both experimentally and theoretically, even for simple lipid mixtures. On the computational side, we have argued that enhanced-sampling all-atom molecular dynamics simulations are uniquely suited for quantification of membrane conformational energetics, not only because they minimize a-priori assumptions, but also because they permit analysis of bilayers in deformed states. To showcase this approach, we examine reported inconsistencies between alternative experimental measurements of bending moduli for cholesterol-enriched membranes. Specifically, we analyze lipid bilayers with different chain saturation, and compute free-energy landscapes for curvature deformations distributed over areas from ∼5 to ∼60 nm^2^. These enhanced simulations, totaling over 100 microseconds of sampling time, enable us to directly quantify both bending and tilt moduli, and to dissect the contributing factors and molecular mechanisms of curvature generation at each length scale. Our results show that cholesterol effects are lipid-specific, in agreement with giant-vesicle measurements, and explain why experiments probing nanometer scale lipid dynamics diverge. In summary, we demonstrate that quantitative structure-mechanics relationships can now be established for heterogenous membranes, paving the way for addressing open fundamental questions in cell membrane mechanics.

**Significance:** Elucidating the energetics and mechanisms of membrane remodeling is an essential step towards understanding cell physiology. This problem is challenging, however, because membrane bending involves both large-scale and atomic-level dynamics, which are difficult to measure simultaneously. A recent controversy regarding the stiffening effect of cholesterol, which is ubiquitous in animal cells, illustrates this challenge. We show how enhanced molecular-dynamics simulations can bridge this length-scale gap and reconcile seemingly incongruent observations. This approach facilitates a conceptual connection between lipid chemistry and membrane mechanics, thereby providing a solid basis for future research on remodeling phenomena, such as in membrane trafficking or viral infection.

## Introduction

All biological membranes resist deformations of their intrinsic shape. A membrane-bound protein may however reshape the surrounding bilayer, sometimes strikingly, because the free-energy cost of membrane bending can be offset by free-energy gains resulting from adequate solvation of the protein surface (see [1] and references therein). In other words, lipid bilayers change shape around proteins to avoid large energetic penalties due to dehydration of ionized surface residues and/or exposure of large hydrophobic clusters to water [2, 3, 4, 5, 6, 7, 8]. This kind of balance between competing energetic contributions is not uncommon, and governs many other processes in molecular membrane physiology, including ligand-induced allostery, ion permeation, and so on. However, while atomic-resolution perspectives have become the state-of-the art in experimental and theoretical analyses of protein structural dynamics, molecular recognition, or solvation energetics, membrane morphology is still often conceptualized assuming a homogeneous medium. While insightful in some cases [9, 10, 11, 12], the shortcomings of this perspective become quickly apparent for mixed-lipid bilayers, which are the norm in biological cells [13]. And so, deceptively simple fundamental questions, such as how the chemical makeup of a bilayer dictates its intrinsic bending energetics, remain largely unresolved.

A recent controversy regarding the stiffening effect of cholesterol illustrates this challenge [14, 15, 16]. Cholesterol is known to be enriched in animal cell membranes, and had been thought to universally enhance their rigidity following the observation that cholesterol dampens the dynamics of the surrounding lipids [17, 18]. If so, generating curvature in cellular membranes would be more costly than in the synthetic bilayers often examined in laboratory conditions, and potentially entail distinct mechanisms. However, while analysis of bilayers of partially or fully saturated lipids confirmed the expected stiffening effect of cholesterol [19, 20], little or no effect was observed for unsaturated lipids like DOPC, when examined either with X-ray diffraction [20], tube aspiration [21, 22] or electro-deformation [23]. This lipid-type specificity would have very interesting biological implications. For example, unsaturated lipids might localize at the periphery of domains enriched in saturated lipids [24] to mitigate the stiffening effect of cholesterol and help preserve the membrane’s plasticity wherever needed. While appealing, however, this lipid-type specificity has been recently disputed by a study of microscopic membrane fluctuations based on neutron-spin echo (NSE), nuclear magnetic resonance (NMR) and conventional molecular dynamics (MD) simulations [14]. In stark contrast to the preceding studies [20, 21, 22, 23], this analysis concluded that the stiffening effect of cholesterol is likely universal for all lipid bilayers, including DOPC, and this notion quickly became the subject of debate [15, 16].

It could be argued that in no small part this controversy stems from the fact that bending moduli are not measured or computed directly, but inferred from other quantities. This route can be misleading when the models used to establish this inference neglect the complex nature of the lipid bilayer, or examine its structure and dynamics without an actual morphological perturbation. In this work, we showcase an advanced all-atom molecular simulation methodology we recently developed [25] to address this kind of challenge. This methodology, which we refer to as ‘Multi-Map’ sampling, provides a means to directly quantify free-energies of bending for lipid bilayers [5], and can be applied to bilayers of any chemical makeup, including model synthetic bilayers with or without cholesterol [25]. This method differs from conventional simulation methods in that it mimics the action of proteins or laboratory manipulations that sustain curvature over long time-scales, while providing atomically-resolved insights into the structure and dynamics of the membrane as it is deformed. Our results unequivocally support the notion that the stiffening effect of cholesterol is nonuniversal [20, 21, 22, 23], and that it is specifically dependent on the degree of lipid-chain unsaturation. Importantly, our results also reveal the mechanism by which lipid unsaturation preserves flexibility despite the increased density and molecular order that cholesterol invariably induces. These insights permit us to explain the divergence between the abovementioned studies of unsaturated DOPC bilayers. In conclusion, we posit that this emerging computational methodology, in combination with suitable experimental approaches, will facilitate a clearer understanding of the molecular mechanisms by which cells and organelles shape the morphology of lipid membranes.

## Results

### Bending free-energy profiles show cholesterol effects are lipid-type specific

Atomistic models of hydrated bilayers were prepared for the following compositions: pure POPC, pure DOPC, and 70:30 mixtures of either POPC or DOPC with cholesterol (CHOL), hereafter indicated as POPC/CHOL and DOPC/CHOL. POPC and DOPC lipids were chosen because they have very similar bending moduli and radii of preferred curvature [26, 27]. A 30% cholesterol mole fraction represents typical concentrations in eukaryotic membranes [28], and approaches the highest concentration that is still soluble in physiological phospholipids [29, 30]. This concentration has also been shown to induce significant changes in molecular properties [14] and membrane mechanics [19, 20].

In order to examine deformations in curvature over different length scales, periodic bilayers of 200, 800 and 1800 lipid molecules per unit cell were prepared for each lipid composition, for a total of 12 membrane models. Respectively, the in-plane dimensions of these molecular systems are approximately 80×80, 160×160 and 240×240 Å ^2^ for the pure bilayers, and 70×70, 140×140 and 210×210 Å ^2^ for the mixed bilayers. All bilayers were simulated in atomistic detail and with explicit solvent at room temperature and pressure using two different sampling methods, generating in each case trajectories of multiple microseconds (Table 1).

**Table 1:** Simulation lengths of all-atom lipid bilayers used in this work. The calculations based on enhanced-sampling simulations entailed multiple replicas with increasing degrees of applied curvature, carried out in parallel.

| Composition | Conventional MD<br>sampling time<br>(single replica), μs | | | Multi-Map enhanced MD<br>sampling time<br>(replica count × time), μs | | |
| --- | --- | --- | --- | --- | --- | --- |
| POPC | 5.3 | 5.9 | 3.3 | $13 \times 0.3$ | $12 \times 0.6$ | $12 \times 1.5$ |
| POPC/CHOL | 8.9 | 9.1 | 3.4 | $13 \times 0.7$ | $14 \times 0.7$ | $13 \times 1.5$ |
| DOPC | 2.0 | 6.5 | 3.3 | $15 \times 0.1$ | $14 \times 0.7$ | $13 \times 1.5$ |
| DOPC/CHOL | 6.2 | 7.5 | 3.4 | $12 \times 0.7$ | $15 \times 0.6$ | $13 \times 1.5$ |
| Number of lipids<br>per unit cell | 1800 | 800 | 200 | 1800 | 800 | 200 |

In one approach, we simulated the bilayers at rest, without applying any extrinsic forces, to examine their spontaneous fluctuations. As will be discussed below, this analysis confirmed that cholesterol increases the molecular order of the lipid bilayer, but to a similar extent for both POPC and DOPC (Figs. 1 and 8).

**Figure 1:**
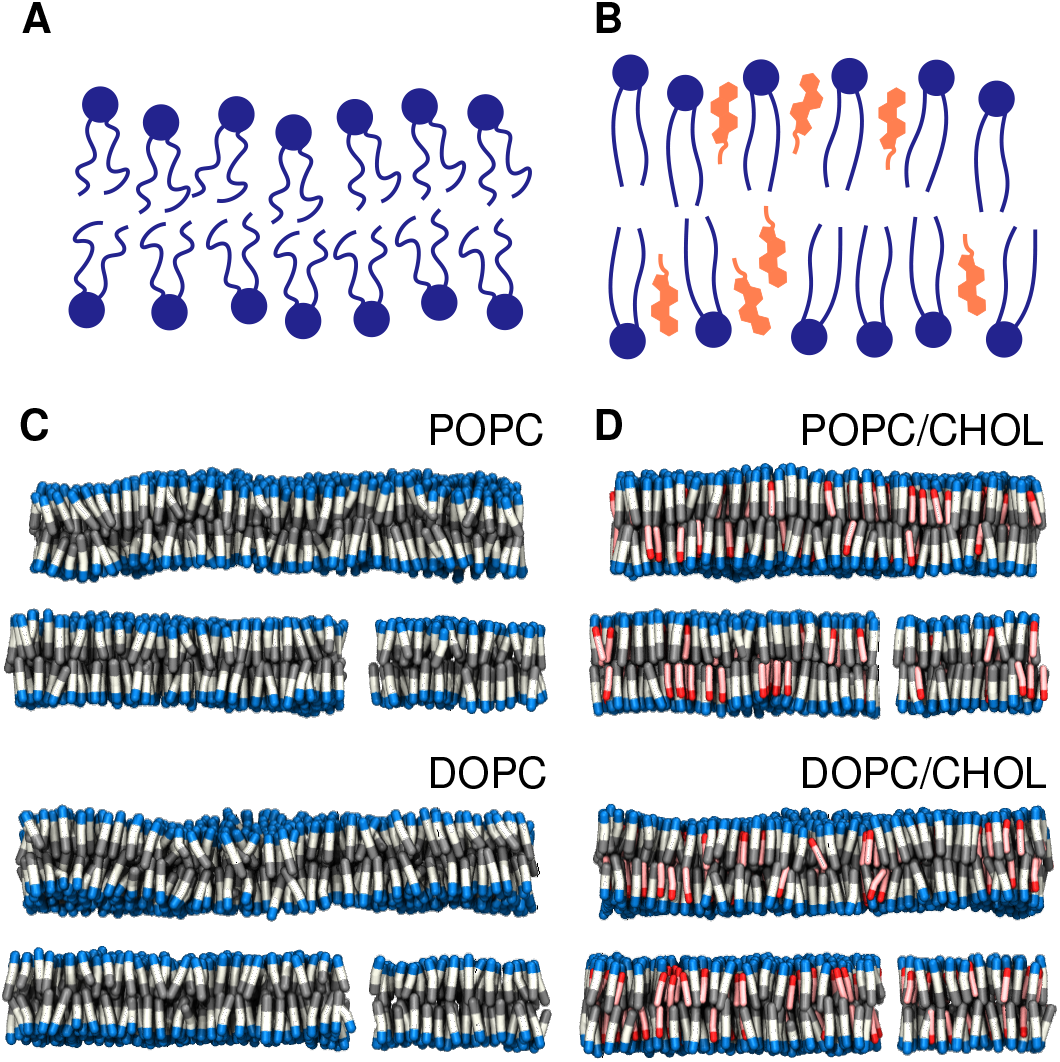
Cholesterol enriched bilayers. **(A, B)** Schematic of the effect of cholesterol on lipid membrane order. **(C)** Snapshots from all-atom simulations of pure-POPC and pure-DOPC bilayers at rest; lipid molecules are visualized as blue/white/gray rods. The water buffer is omitted for clarity. **(D)** Simulation snapshots of POPC/CHOL and DOPC/CHOL bilayers. Cholesterol molecules are represented as red/pink rods.

To more directly evaluate the effect of cholesterol on the energetics of membrane bending, we used the Multi-Map enhanced-sampling method [25]. In this approach, a collective variable (or reaction coordinate) that reflects the membrane shape and curvature [25] is gradually forced to explore a range of values, by applying a series of progressively shifted biasing potentials following the umbrella-sampling protocol [31]. Here, the bilayers were driven to adopt a periodic sinusoidal shape along the *x*-axis, gradually varying the amplitude of the sinusoid to induce greater curvature. It is important to note that while this bias fosters an average shape in the membrane, it does not influence its fluctuations around that shape, nor does it impose one or other mechanism of bending at the molecular level. Through post-processing of these enhanced simulations, we then derived for each bilayer the potential of mean force (PMF), or free-energy profile, as a function of the applied curvature *c*.

These PMF profiles, shown in Fig. 2, clearly demonstrate that bending a POPC/CHOL bilayer is significantly more costly than bending a POPC bilayer, for curvature deformations of comparable shape and length-scale (Fig. 2). That is, cholesterol has a substantial stiffening effect on POPC membranes. By contrast, the free-energy curves for DOPC and DOPC/CHOL are nearly identical (Fig. 2), i.e. cholesterol has a minimal influence on the plasticity of the DOPC membranes, despite its effect on molecular order.

**Figure 2:**
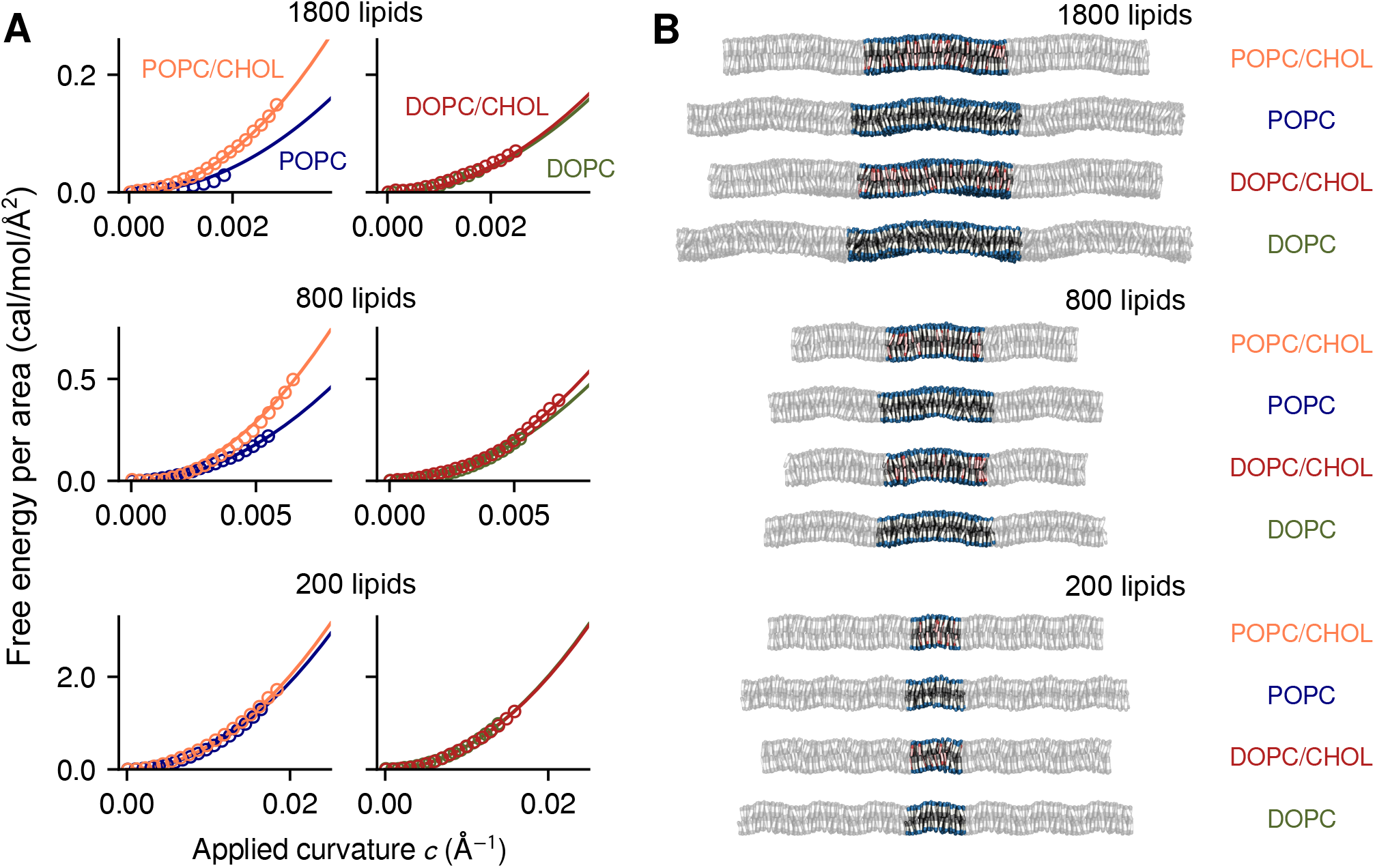
Direct calculation of the free-energy cost of lipid bilayer bending using the Multi-Map enhanced-sampling simulation method. **(A)** Potentials-of-mean-force (PMFs) as a function of bilayer curvature for an applied sinusoidal shape along the x-axis. Data are shown for POPC (*blue*), POPC/CHOL (*orange*), DOPC (*green*) and DOPC/CHOL (*red*) bilayers, containing either 1800, 800 or 200 lipid molecules (as indicated). All PMF profiles are given in units of free-energy per area; symbols indicate computed values, and solid lines are quadratic fits. **(B)** Representative snapshots of each bilayer, represented as in Fig. 1. The figure highlights the periodic unit cell, flanked by its images along the direction of the sinusoid.

It is worth noting that the statistical error of the PMF profiles shown in Fig. 2 is very small, despite the large computational cost of the calculations; thus, these curves are faithful representations of the free-energy cost of membrane bending at the corresponding length scale, given the forcefield used in the simulations. From these data alone, therefore, it is apparent that the stiffening effect of cholesterol is lipid-type specific rather than universal. Nonetheless, to enable direct comparison with other experimental and computational approaches, it is useful to further quantify this stiffening effect through the bending modulus *k*_*c*_. Derivation of *k*_*c*_, however, requires further analysis of our curved-membrane trajectories, because the dominant mechanisms of bending depend on the length scale of the curvature deformation at the molecular level [32, 33, 34, 35, 27, 36]. This problem is addressed in the next section.

### Determination of bending and tilt moduli from analysis of curved lipid bilayers1

The Helfrich- Canham theory [37, 38] idealizes the lipid bilayer as a homogeneous, harmonic elastic medium represented by its midplane. The free-energy density of bending this membrane, *f*_HC_, relies on a single parameter, the bending modulus *k*_*c*_, and has the following expression:

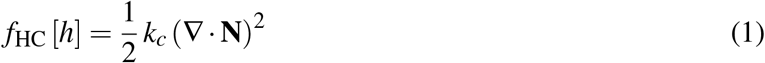

where *h*(*x, y*) is a two-dimensional function that maps the difference between the shape of the deformed bilayer and a flat shape. *f*_HC_ depends directly on the second derivatives of *h*, or equivalently the first derivatives of the vector perpendicular to the membrane midplane **N** = (*∂*_*x*_*h, ∂*_*y*_*h*, −1), save for normalization. The Helfrich-Canham model therefore implicitly assumes a mechanism wherein, on average, all lipid molecules remain parallel to the normal vector **N**, and therefore bending entails deflections in the relative orientations of adjacent lipid molecules (Fig. 3A).

**Figure 3:**
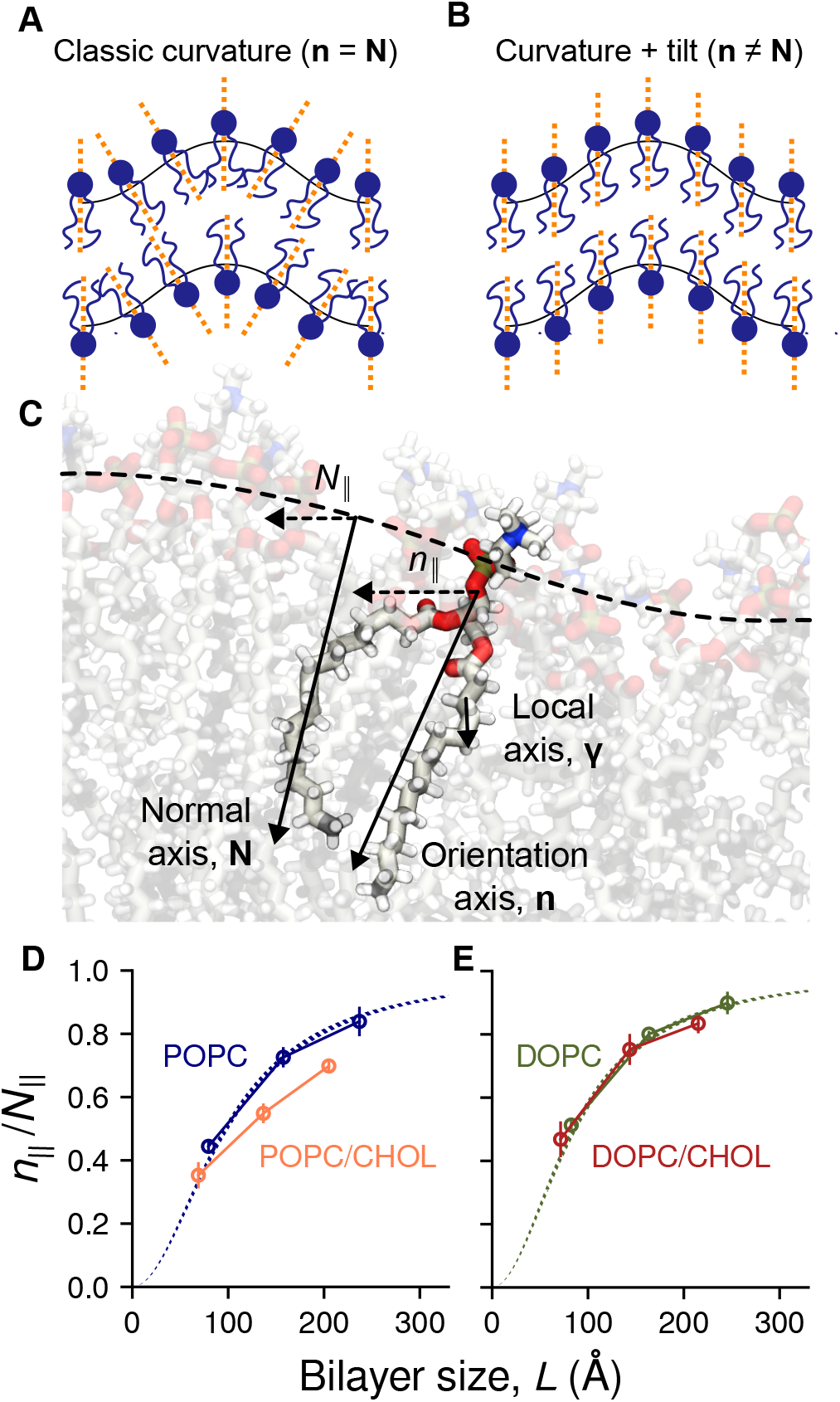
Membrane bending entails midplane curvature and lipid tilt. **(A)** Schematic of an idealized curved bilayer wherein one or other term of Eq. 2 [32] dominates the energetics. **(B)** Illustration of the membrane normal vector, **N**, the lipid orientation vector (or lipid director), **n**, and the local chain vector *γ*, analyzed in Fig. 6). The ratio *n*_∥_*/N*_∥_ measures the degree to which midplane curvature defines the bending energetics; *n*_∥_*/N*_∥_ = 1 only in the ‘macroscopic’ limit, where the Helfrich-Canham model is valid. **(C)** Mean values of *n*_∥_*/N*_∥_ for POPC (*blue*), POPC/CHOL (*orange*), DOPC (*green*) and DOPC/CHOL (*red*), calculated from the same MD trajectories used to derive the PMF profiles in Fig. 2. Error bars represent standard errors. Dashed lines show the theoretical predictions for DOPC and POPC using Eq. 3 and published values of *k*_*c*_*/k*_*t*_ [26].

While this assumption is plausible for deformations occurring over ‘macroscopic’ length-scales, it is increasingly recognized that the average lipid orientation in more localized bending is not solely described by the shape of the membrane midplane; that is, lipids can ‘tilt’ relative to the normal vector **N** (Fig. 3B). A more general free-energy functional is therefore the following [32, 33]:

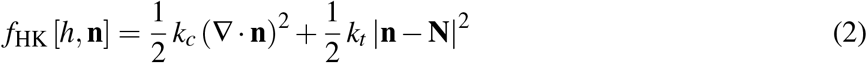

where **n** maps the lipid orientation across the membrane and *k*_*t*_ is referred to as the lipid-tilt modulus. Note that while the first term in Eq. 2 is very similar to Eq. 1, **n** varies more smoothly than **N** at short length scales (*<* 100 Å, Fig. 3B) and thus the free-energy cost of bending according to Eq. 2 can be significantly smaller than that predicted by the Helfrich-Canham model (Eq. 1). The spectral density of fluctuations generated by Eq. 2 has been supported by many simulations and experiments [39, 40, 34, 27, 36], but to estimate *k*_*c*_ and *k*_*t*_ can prove challenging, particularly for non-homogenous membranes.

A common approach to estimate *k*_*c*_ and *k*_*t*_ is to examine the spontaneous fluctuations of a membrane at rest, either by simulation or experiment. However, because membrane fluctuations are transient, **n** and **N** are difficult to resolve in both position and time, requiring substantial averaging. This challenge is greater when the intrinsic bilayer shape is flat, as **n** and **N** fluctuate around the same fixed direction, and only their fluctuations carry physical information. By contrast, evaluation of **n** and **N** is more accurate when the membrane is examined under applied curvature, as each element varies very significantly across the bilayer, to an extent that can be predicted by theory.

For example, for a periodic sinusoidal shape along the *x*-axis, it is straightforward to evaluate the three components of **N** and their derivatives, since the only longitudinal component is *N*_*x*_, hereafter labeled as *N*_∥_. By minimizing Eq. 2 with respect to the corresponding orientation component, *n*_∥_, we obtain the following expression:

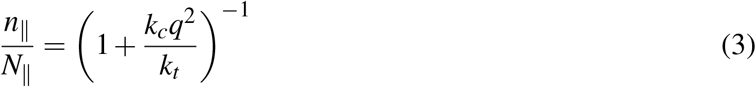

where *q* = 2*π/L* and *L* is the length of the sinusoidal curvature deformation (i.e. the bilayer size). Note that because *N* and *n* are constant along the *y* axis, the “twist” term ∇ × (**n** − **N**) [32, 34, 36] can be safely neglected and thus was not included here for clarity. Splay-tilt coupling [41] was similarly neglected because it is thought to be significant only at very high *q* [36].

Based on the same trajectory data used for the derivation of the PMF curves (Fig. 2) we calculated the ratio *n*_∥_*/N*_∥_ for each bilayer using linear regression (Fig. S1), consistent with the underlying assumption of linear response [32]. For pure POPC and DOPC we find that the simulation data and Eq. 3 are in excellent agreement when using known values of *k*_*c*_*/k*_*t*_ [26] without any additional fitting (Fig. 3D and E). This is a non-trivial result that confirms our methodology based on analysis of extrinsically-curved bilayers is sound. We thus proceeded to determine *k*_*c*_ and *k*_*t*_ for POPC/CHOL and DOPC/CHOL membranes, which to our knowledge had not been previously estimated. We again find that Eq. 3 also describes our data very well for these mixtures, using the *k*_*c*_*/k*_*t*_ ratio as the only fitting parameter (Fig. S2).

The results of this analysis enable us to dissect the effect of cholesterol on bending and tilting for each lipid type. As shown in Fig. 4 and Table 2, we find that *k*_*c*_ increases by a factor of 2.1 *±* 0.2 when adding cholesterol to POPC (Fig. 4A), confirming that cholesterol has a significant stiffening effect for this type of lipid. Our estimate is in excellent agreement with tube-aspiration measurements, which yield a 2.3-fold increase [19]. By contrast, the relative change in *k*_*c*_ between DOPC and DOPC/CHOL is 1.1 *±* 0.1 (Fig. 4B), consistent with multiple measurements that indicate no stiffening [20, 21, 22, 23]. Although those experiments probed “macroscopic” length-scales, our results now show that the effect of cholesterol on DOPC is also minimal at length scales comparable to the membrane thickness (*L <* 100 Å). Indeed, the effect of cholesterol in this scale is also small for POPC (see 200-lipid system, Fig. 2, as tilting becomes more dominant, thereby mitigating the free-energy cost of curvature (Fig. 3D).

**Table 2:** Bending and tilt moduli for pure and mixed bilayers (means and 95% confidence intervals over the three bilayer sizes).

| | $k_c$ (kcal/mol) | $k_t$ (kcal/mol/Å<sup>2</sup>) |
| --- | --- | --- |
| POPC | 21.6 [17.6, 25.6] | 0.104 [0.085, 0.124] |
| POPC/CHOL | 45.7 [41.2, 50.2] | 0.111 [0.100, 0.122] |
| DOPC | 20.9 [17.1, 24.7] | 0.114 [0.094, 0.134] |
| DOPC/CHOL | 23.8 [21.5, 26.1] | 0.134 [0.121, 0.147] |

**Figure 4:**
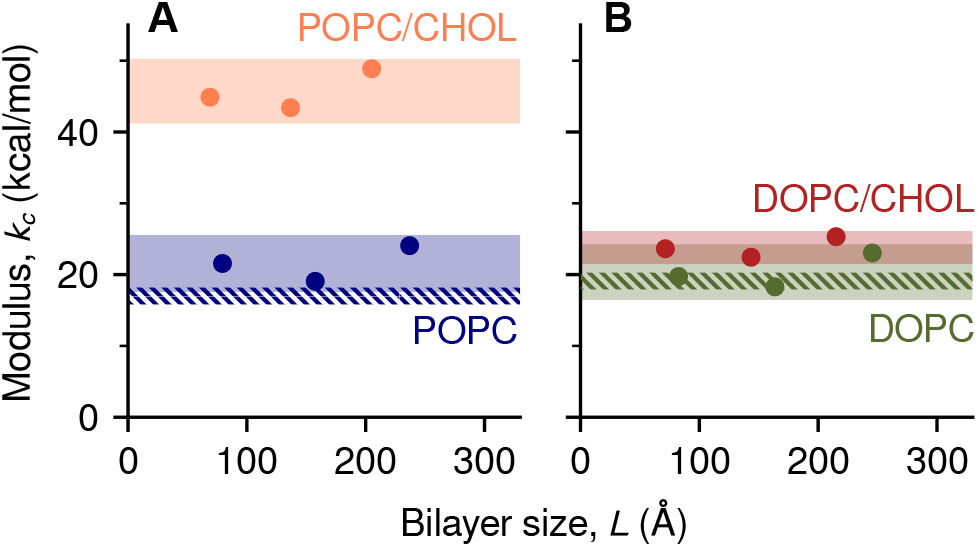
The stiffening effect of cholesterol is lipid-type specific. Calculated bending moduli kc are plotted as a function of bilayer size for different bilayers of **(A)** POPC (*orange*) and POPC/CHOL (*blue*), and **(B)** DOPC (*green*) and DOPC/CHOL (*red*). All values were derived from analysis of MD trajectories of curved bilayers, as in Fig. 2. Shaded bands indicate the estimated 95% CI of *k*_*c*_ (Table 2, and dashed bands indicate the 95% CIs of *k*_*c*_ from tilt fluctuations of pure lipids [26].

In summary, our simulation data unequivocally show that the stiffening effect of cholesterol is nonuniversal. More specifically, stiffening is very pronounced for POPC but marginal for an unsaturated lipid like DOPC. It is worth noting that the effect of cholesterol on POPC bilayers can also be small, but only for very localized deformations. At larger length scales, however, cholesterol causes POPC and DOPC bilayers to have a very different plasticity.

### Rotational dynamics around unsaturated bonds explains differential cholesterol effect

Although Eq. 2 may be used to calculate the free-energy of membrane bending when empirical parameters such as *k*_*c*_ and *k*_*t*_ are known or assumed, no continuum theory can explain how these parameters depend (or not) on the lipid composition of the membrane. This explanation can only be obtained by examining the internal molecular structure and dynamics of curved bilayers and the precise configuration of their constituent lipids. To that end, we used our trajectory data to examine several plausible descriptors, namely membrane thickness and local cholesterol enrichment, as well as hydration of the hydrocarbon chains and their rotational freedom. All these properties fluctuate on time scales shorter than microseconds, and therefore our trajectories yield excellent statistics.

We observed no significant correlations between applied curvature and local bilayer thickness for either lipid type or length scale (Fig. 5A and B), ruling out this property as a reason for the differential effect of cholesterol. For the more localized deformations (*L <* 100 Å), we observed a small increase in the local cholesterol concentration (*<* 0.5 mol%) where the membrane has the highest curvature (*c ≈* 0.02 Å ^−1^, Fig. 5C,D). However, this modest enrichment is similarly small for both POPC/CHOL and DOPC/CHOL. Therefore, local enrichment does not explain either why cholesterol stiffens POPC but not DOPC (Fig. 2); instead, this enrichment likely reflects a general preference of cholesterol for concave curvatures.

**Figure 5:**
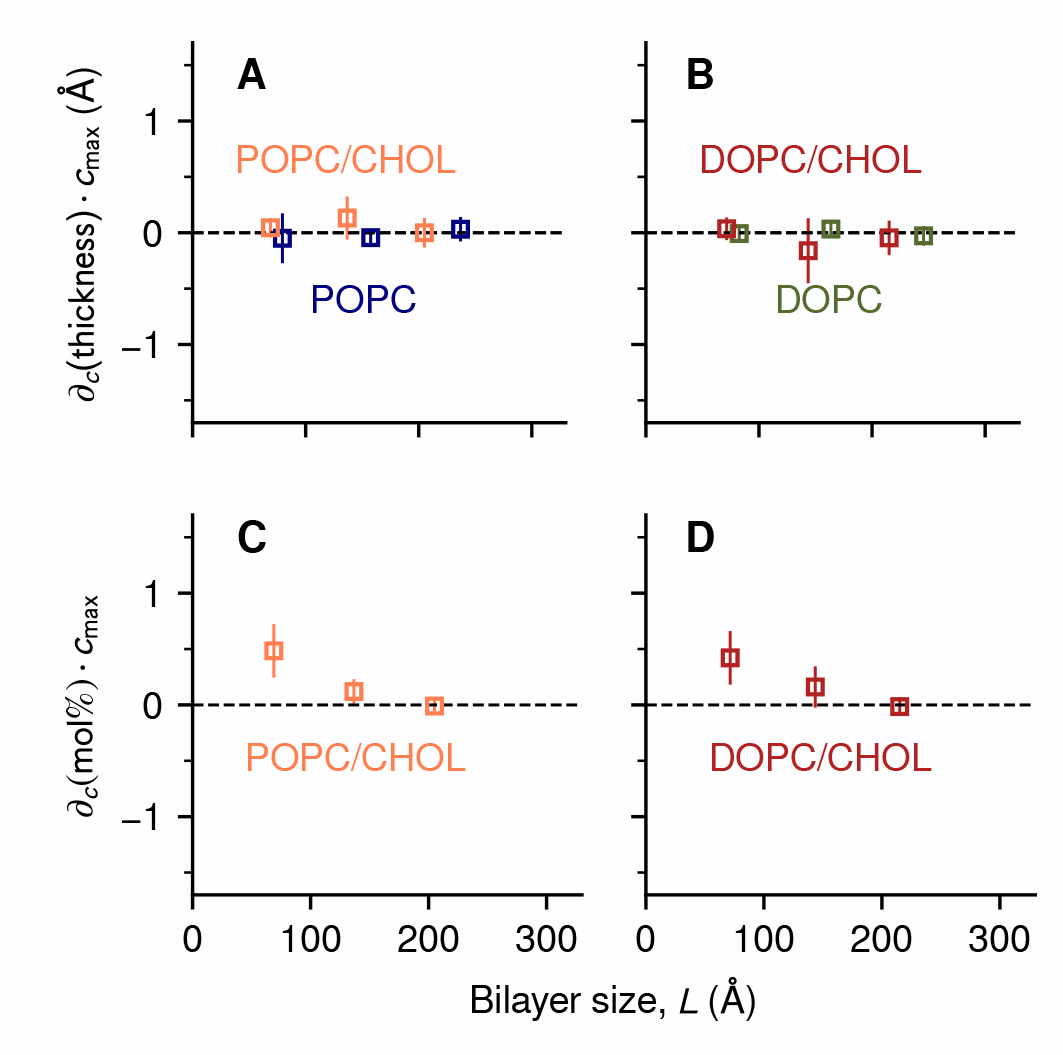
Differential stiffening effect is not explained by curvature-induced variations in bilayer thickness or local cholesterol enrichment. **(A, B)** Derivatives of the thickness with respect to curvature for different bilayers of **(A)** POPC (*blue*) and POPC/CHOL (*orange*), or **(B)** DOPC (*green*) and DOPC/CHOL (*red*). The derivatives were estimated by linear regression; for ease of comparison, the resulting slopes were multiplied by the largest applied curvature *c*_max_. **(C, D)** Derivatives of the cholesterol concentration with respect to curvature for different bilayers of **(C)** POPC/CHOL (*orange*) and DOPC/CHOL (*red*). Error bars represent standard errors.

Analyzing the hydration of the hydrocarbon chains, we found a clear correlation between curvature and water exposure of the acyl chains, as expected, but only for the pure-lipid bilayers and not for their cholesterol mixtures (Fig. S3), therefore ruling out this factor.

The differential effect of cholesterol on DOPC and POPC bilayers was however revealed when examining the extent to which increasing curvature alters the orientational dynamics of individual acyl-chain segments. This analysis entails evaluation of the vectors formed by every pair of carbon atoms spaced by two chemical bonds in each lipid chain:

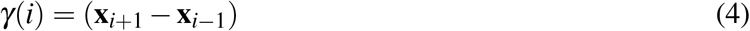

Similar to the analysis associated with Eq. 3, the longitudinal component of these vectors, *γ*_∥_ (*i*) (Fig. 3C), was evaluated and the ratio *γ*_∥_ (*i*)*/N*_∥_ quantified through regression. The result was then normalized against the relative orientation of the entire lipid (*n*_*>∥*_*/N*_*>∥*_), to quantify the dynamics of the *i*-th segment relative to other segments in the same lipid chain. A higher value of *γ*_*>∥*_(*i*)*/N*_*>∥*_ indicates a greater deflection in that particular chain segment in response to bilayer curvature.

The results of this analysis are shown in Fig. 6. In the absence of cholesterol, *γ* _*>∥*_(*i*)*/N*_*>∥*_ is highest for the chain segments that are most proximal to the head groups (*i <* 3) and decays rapidly over a distance shorter than the chain persistence length (*≈* 6 Å). Cholesterol lowers the value of *γ* _*>∥*_(*i*)*/N*_*>∥*_ even further, with the largest effects observed in the saturated *sn*-1 chain of POPC (Fig. 6A), and near the sterol group in general (4 ≤ *i* ≤ 10). It is outside of this region, however, that DOPC and POPC respond to curvature deformations in entirely different ways, which in turn explains the distinct effect of cholesterol.

**Figure 6:**
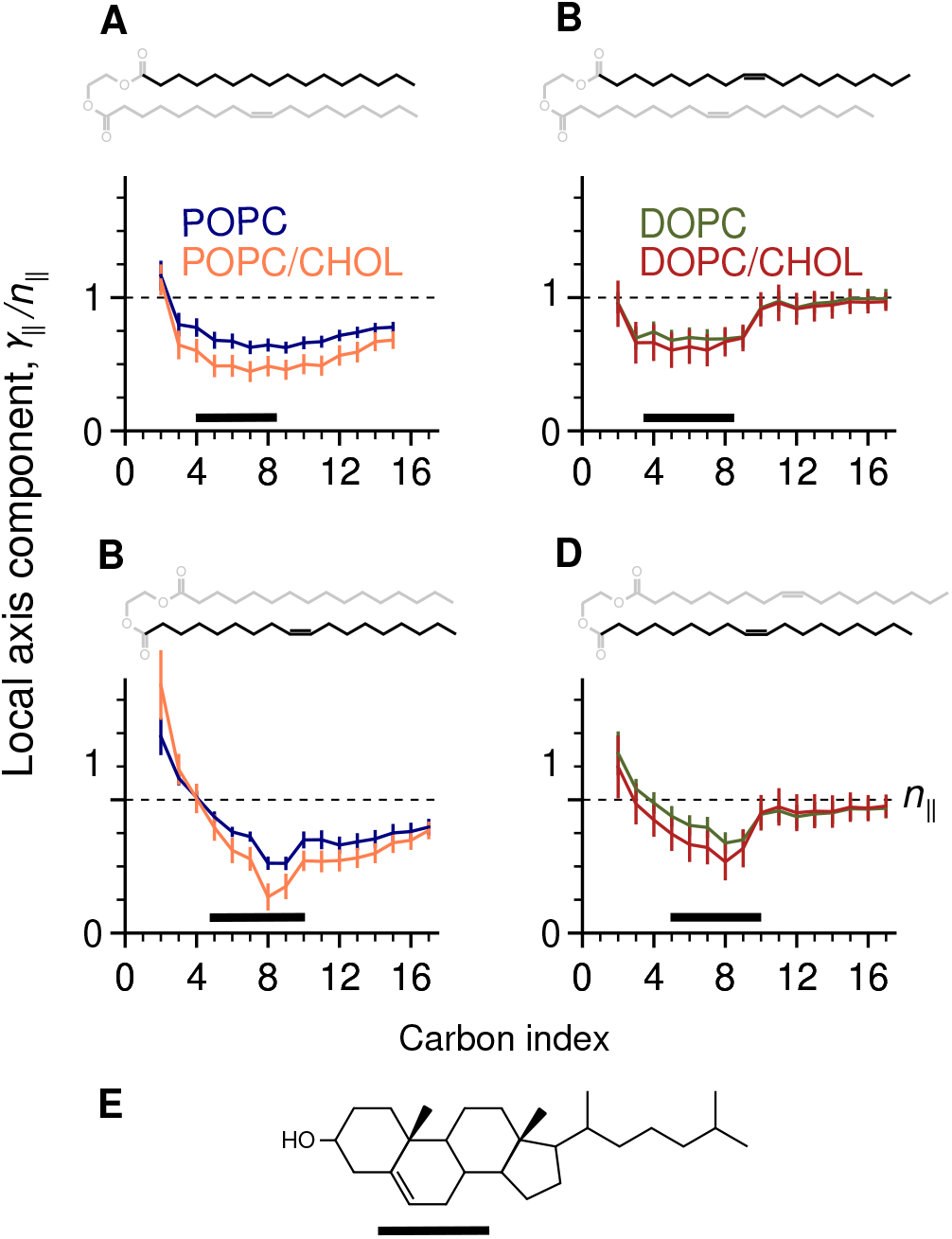
Concurrent and strategically located unsaturated bonds confer the ability to adapt to curvature. Shown are mean values of the longitudinal component *γ*_∥_ of the local chain axis (Fig. 3C), normalized against the lipid molecule’s orientation *n*_∥_, shown as dashed lines. **(A, B)** Values of *γ*_∥_ acyl chain segments in POPC (*blue*) and POPC/CHOL (*orange*), focused on either **(A)** the *sn*-1 chain or **(B)** *sn*-2 chain. **(C, D)** Equivalent results for DOPC (*green*) and DOPC/CHOL (*red*). In each panel the chemical structure of the acyl chain is highlighted. Black bars indicate the central 50% of the density of the sterol group, shown in **(E)** for reference.

In DOPC, segments beyond the 9^th^ carbon in each chain pivot relatively freely, manifested as *γ* _*>∥*_ *≃ n*_*>∥*_, and the “hinge points” of this mobility are found at the unsaturated bonds (Fig. 6C,D). Strikingly, the ability of these segments to move is completely unaffected by cholesterol, explaining why the free-energy of membrane bending does not increase for DOPC/CHOL.

This rotational dynamics is entirely different for POPC: the unsaturated *sn*-2 chain has little effect on the saturated *sn*-1 chain (Fig. 6A), which remains relatively rigid and shows limited ability to rotate. When cholesterol is added, the rigidity of the *sn*-1 chain increases further, explaining the higher free-energy cost needed to generate the same curvature.

### Analysis of spontaneous fluctuations of membranes at rest is inconclusive

In the preceding sections we have shown how specialized MD simulations designed to induce steady-state curvature deformations in membranes can be used not only to quantify their bending energetics and empirical parameters often used in analytical theories, but also to pinpoint the molecular origins of observed variations against lipid composition. Studies of lipid bilayers at rest using conventional MD simulations are however far more common; in this section, we show that while analysis of spontaneous fluctuations is certainly insightful, the information gained by this route is ultimately limited.

Specifically, for each of the lipid bilayers considered above, now examined at rest, we calculated an MD trajectory spanning multiple microseconds (Table 1). We then quantified the spontaneous shape fluctuations of the membrane midplane for each trajectory snapshot by evaluating the instantaneous position of each lipid molecule (represented by its center of mass) relative to the bilayer center, i.e. *h*(*x*_*i*_, *y*_*i*_) = (*z*_*i*_ − ⟨*z*⟩). The 2D Fourier transform of *h*, denoted by *h*_**q**_, was then computed over all wave vectors with wavenumber |**q**| *<* 0.2 Å ^−1^, which is the appropriate limit for all-atom simulations [36]. The resulting values of |*h*_**q**_|^2^ were then averaged over all trajectory snapshots collected for *t >* 0.5 *μ*s, and data points with equal |**q**| combined. To relate the calculated spectrum of bending fluctuations to the bending and tilt moduli, we used the following expression, based on Eq. 2 [32, 34, 27, 36]:

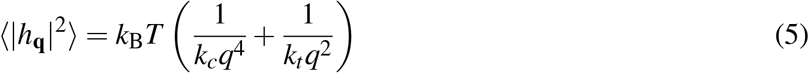

The results of this analysis are shown in Fig. 7 and Table S1. We find that the bending moduli *k*_*c*_ can be determined with reasonable certainty for both POPC and DOPC membranes (∼15% error); the estimated values agree well with those obtained with previous MD simulations or “flicker” spectroscopy [34, 26], though they are slightly larger than those deduced from X-ray diffraction data [27]. (This discrepancy has been discussed elsewhere and remains to be understood [26, 27]: it has been speculated that the close spacing of the multi-lamellar stacks used for X-ray diffraction might introduce a favorable coupling between bilayers [26].) Nonetheless, all methods agree that POPC and DOPC bilayers have nearly identical *k*_*c*_ and *k*_*t*_.

**Figure 7:**
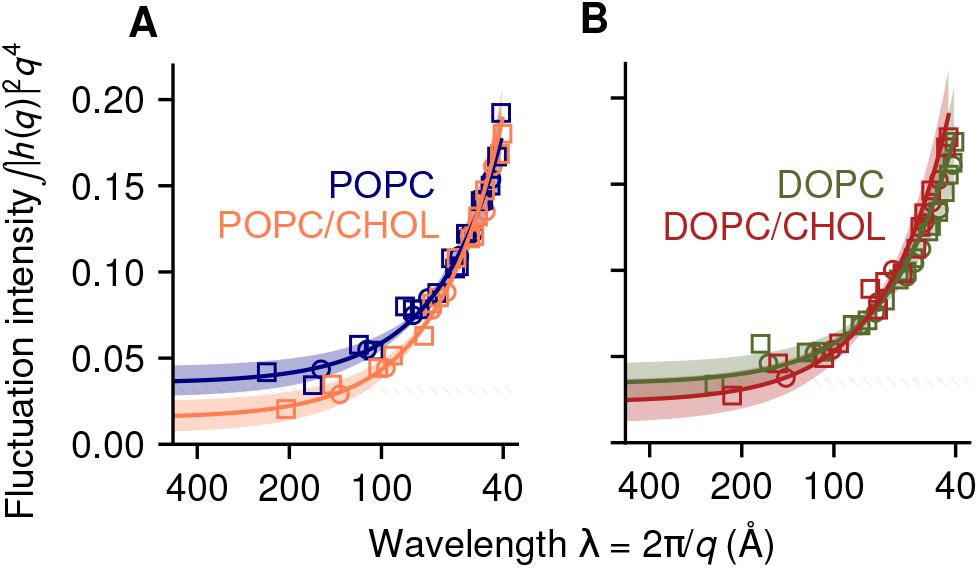
Spectral analysis of bending fluctuations of bilayers at rest, based on conventional MD simulations. **(A)** POPC and POPC/CHOL bilayers (*\* * P* = 0.002). **(B)** DOPC and DOPC/CHOL bilayers (*P* = 0.24). Solid lines are best fits of the function in Eq. 5, with shaded bands indicating the 95% confidence interval (CI). *P*-values are for comparisons of *k*_*c*_ with and without cholesterol. For comparison, dashed bands show the 95% CI for POPC and DOPC under the *q*^−4^ long-wavelength Helfrich-Canham theory [37, 38], using published values of *k*_*c*_ [26].

**Figure 8:**
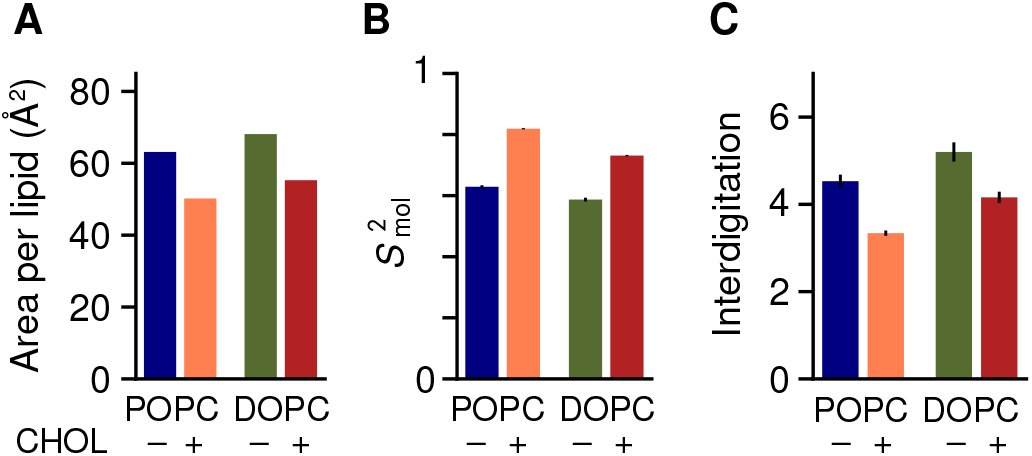
Non-specific cholesterol effects on membrane order and structure, irrespective of lipid-chain saturation. **(A)** Area per phospholipid molecule computed for bilayers of POPC (*blue*), POPC/CHOL (*orange*), DOPC (*green*) and DOPC/CHOL (*red*). **(B)** Orientational order parameter 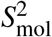 of lipid molecules, for the same bilayers. **(C)** Degree of entanglement between leaflets, estimated using the number of interdigitating acyl-chain bonds.

For POPC/CHOL and DOPC/CHOL, the statistical uncertainties are much higher (∼35%, Table S1); as noted above, analysis of membranes at rest is more prone to sampling deficiencies, and this problem is exacerbated when molecular motions are dampened, as in the presence of cholesterol. Nonetheless, we can resolve a significant difference between the bending moduli *k*_*c*_ of POPC and POPC/CHOL (\*\**P* = 0.002; Fig. 7), consistent with the 2.3-fold increase measured by tube aspiration [19]. However, it was not possible to detect significant changes between DOPC and DOPC/CHOL; in the light of our direct analysis of bending energetics, little or no difference is indeed the expected result, but without that information, the uncertainty inherent to this type of analysis would preclude a conclusive quantitative answer to the question at hand.

For completeness, we note that the spontaneous fluctuations of a membrane at rest might be examined differently; for example, a method has been reported that analyzes the fluctuations of the lipid-tilt field (**n** − **N**), which are uniquely accessible to MD simulation [34]. (NMR also measures the fluctuations of **n**, but they are aggregated with those of **N** as well as movements of individual atoms [42].) While this tilt-fluctuations analysis [34] can be more precise than the bending-fluctuations analysis (Eq. 5), the former has not yet been validated on mixed-composition bilayers and therefore we did not consider it here.

### Seemingly universal cholesterol effects

To conclude, it is worth clarifying that while cholesterol might not have a strong influence on the bending energetics of bilayers of a certain lipid-type, some effects do appear to be universal. For example, there is consensus that when cholesterol is mixed with any kind of phospholipids, it significantly dampens their motion and increases their alignment with the membrane normal, as illustrated in Fig. 1A,B. Both effects are indeed observed in our simulations of membranes at rest, and translate into clear differences in the area per lipid molecule (Fig. 8A) and in the atomic order parameters (Fig. S4). Consistent with earlier analyses [43], the difference is higher for saturated chains (*sn*-1 chain of POPC) than for unsaturated chains (DOPC and *sn*-2 chain of POPC). An alternative metric of chain entropy that is independent of overall molecular orientation can be obtained by quantification of the rotational freedom of individual torsional angles, independent of overall molecular orientation (Fig. S5). This metric shows that adding cholesterol results in a significant loss of entropy per molecule for both POPC and DOPC, namely −*T* Δ*S* = 1.05 kcal/mol and 0.44 kcal/mol, respectively (Fig. S5).

An additional route to quantify the orientational order of lipid molecules is through the auto-correlation function (ACF) of the lipid orientation vector:

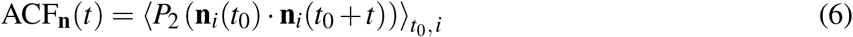

where **n**_*i*_ is the orientation vector of the *i*-th lipid molecule, *t*_0_ and *t* are time intervals, and *P*_2_(*x*) = (3*x*^2^ − 1)*/*2. Lipid orientation fluctuations are much faster than membrane curvature fluctuations [34, 26]: therefore, Eq. 6 has a relatively well defined plateau near *t* = 100 ns, here indicated as 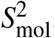. Cholesterol causes a significant increase in 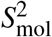 for both POPC and DOPC (Fig. 8B). We note that values of 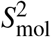 we obtain for DOPC and DOPC/CHOL (0.59 and 0.73, respectively) are in close agreement with those reported elsewhere [14].

## Discussion

To examine opposite claims in regard to the universality of cholesterol stiffening of lipid bilayers, resulting from distinct experimental approaches [15, 16], we used a recently developed simulation methodology [25] to compute bending free-energy curves for atomically-detailed bilayers of POPC, DOPC and their mixtures with 30 mol% cholesterol, at length scales relevant in molecular physiology (∼100-250 Å). Our results clearly show that while POPC and DOPC have virtually identical mechanical properties, the addition of 30 mol% cholesterol increases the bending modulus *k*_*c*_ of POPC over two-fold but has little effect on DOPC. These results are thus in line with a body of experimental evidence that supports the notion that the stiffening effect of cholesterol is lipid-type specific, rather than universal [19, 20, 21, 22, 23]. Evidently, this conclusion rests on two key elements of our theoretical analysis, namely the force field used in the simulations [44], and the curvature-tilt energy functional described in Eq. 2 [32]. The validity of both elements is however well documented [26, 39, 40, 34, 27, 36], and so it is difficult to envisage that the opposite conclusion in regard to the stiffening effect of cholesterol could emerge from an alternative analysis of simulation data.

Because our computational methodology entails simulation of all-atom trajectories, it permits us to not only quantify the bending energetics of the membrane but also to inspect its origins with molecular resolution. Our trajectory data show that while POPC and DOPC bilayers appear comparable when described by macroscopic parameters such as bending modulus or preferred curvature, they differ significantly when examined at the atomic level under applied curvature, and these differences in turn explain why cholesterol has a distinct effect in each case. Specifically, we observe that the unsaturated bonds cause both chains in DOPC to change orientation much more dynamically than the chains in POPC; this configurational freedom is not impacted for DOPC/CHOL owing to the precise location of these bonds relative to the preferred location of cholesterol in the membrane (Fig. 6). Interestingly, this location is highly conserved in mono-unsaturated lipid chains [45, 43], suggesting that this physical property serves important biological functions.

Our results for DOPC and DOPC/CHOL are at odds with the conclusions of a recent study based on NMR and NSE measurements, as well as conventional MD simulations, which proposed that cholesterol also stiffens DOPC membranes [14]. It is important to note, however, that this conclusion is inferred from an evaluation of molecular-scale spontaneous fluctuations at rest, and not from a direct quantification of the free-energy cost of membrane bending; as is common, this inference hinges upon specific theoretical models and approximations. For example, Chakraborty *et al* [14] used indirectly the Helfrich-Canham equation [37, 38] to parameterize the membrane bending energetics. Although effective in the ‘macroscopic’ regime, this theory is inadequate at the nanometer scale [39, 40, 34, 27, 36]. The inadequacy of this approximation is clearly illustrated in Fig. 7, which demonstrates that the magnitude of the spontaneous membrane bending fluctuations significantly exceeds the Helfrich-Canham prediction, to an extent that depends both on length scale and lipid composition.

It is also worth noting that the NSE method relies on experimental detection of the relaxation rates of collective membrane fluctuations in the nanometer range and over hundreds of nanoseconds [46, 14]. This approach can thus yield unique insights into the structural dynamics of a mixed membrane; for example, NSE measurements have recently demonstrated that binary mixtures of saturated lipids (DMPC/DSPC) have nonadditive properties [47]. However, because NSE is sensitive to different kinds of membrane fluctuations, theoretical models are used to isolate each contribution when interpreting NSE data. Specifically, bending fluctuations are estimated by analyzing NSE-measured decay rates as a function of wavelength [48, 49]. Although the underlying theory acknowledges the existence of “fast” and “slow” exponential decays [50], single exponentials are assumed in that analysis due to the short time span of NSE experiments [49]. We tested this assumption by examining the time auto-correlation functions of the Fourier coefficients *h*_**q**_, whose variances |*h*_**q**_|^2^ were previously analyzed in Fig. 7. Importantly, we found that *h*_**q**_ decays as a single exponential only for pure POPC or pure DOPC, but not for POPC/CHOL and DOPC/CHOL (Fig. S6A-H). The reason for this deviation is unclear and deserves further examination; at any rate, we would argue that this finding calls for a reassessment of the model used to extract bending moduli from NSE data [14], e.g. by accounting explicitly for multiple dissipative motions in the membrane. Because interleaflet friction can now be measured directly with reasonable accuracy [51], future NSE studies might leverage other experiments to reduce the number of theoretical assumptions.

Unlike NSE, NMR experiments examine membrane dynamics by measuring the orientational order of specific chemical bonds in the lipid chains. Atomic, molecular and collective fluctuations all contribute together, and are analyzed using a kinetic theory that correlates order parameters with relaxation rates for each bond [42]. Specifically, extraction of a bending modulus *k*_*c*_ from NMR data involves synthesis of all collective membrane fluctuations through a fixed “slow” order parameter, *S*_s_ [42, 14, 52]. A caveat of this approximation is therefore that the relative magnitudes of the different types of fluctuations are implicitly imposed by the model, rather than being specifically measured for each membrane.

Lastly, while MD simulations of bilayers at rest in principle offer the means to quantify each type of fluctuation independently, statistical sampling of spontaneous bending motions (i.e. collective) is often too limited, both in length and time scale [36]. Given these limitations, bending rigidity is typically inferred from analysis of single-molecule descriptors. For example, Chakrarborty *et al* [14] examine the fluctuations of the “splay” angle between pairs of neighboring lipid molecules. We reproduce those results in Fig. S7, but question this route to derive macroscopic moduli. One of the underlying assumptions of this approach is that the fluctuations of the lipid tilt vector (**n** − **N**) are decoupled from those of the normal vector, **N** [53]. This approximation is reasonable only in the ‘macroscopic’ regime, and requires adopting a moving frame of reference to account for the fluctuations of **N**, without assuming it constant over the bilayer [34, 26]. This problem is innate in the analysis of membrane fluctuations at rest, which we avoided by examining changes in **n** and **N** that significantly exceed those fluctuations, in deformed membranes.

We conclude this discussion by underscoring a key result specific to the shortest length scales considered in our simulations, namely *L <* 100 Å. At this length scale, the mechanism of curvature generation remains unchanged upon addition of cholesterol, and there is little or no stiffening effect for *either* DOPC or POPC (Fig. 2). This is an important observation, because it suggests that individual membrane proteins interact with a mechanically invariant environment even when the neighboring concentration of cholesterol is substantial. In other words, while cholesterol can impact membrane remodeling at scales comparable to lipid domains [24], it would not disrupt the activity of individual membrane proteins whose mechanisms entail local deformations of the membrane [5, 54]. Quantitatively testing this hypothesis will prove essential for establishing how protein biochemistry dictates membrane morphology.

## Methods

### Systems, simulation specifications and trajectory analysis

Initial all-atom models of hydrated lipid bilayers were constructed by replicating pre-equilibrated configurations from an earlier study [25]. The distance separating periodic images of each bilayer along the perpendicular to the midplane is about 70 Å ; this water content is about four times greater than in multil-amellar samples used for X-ray diffraction [27]. Interatomic forces were represented via the CHARMM36 and TIP3P force fields [44, 55] using a 12-Å cut-off for Lennard-Jones forces and constraints for all covalent bonds involving hydrogen atoms. All simulations were performed with NAMD [56] at 300 K and 1 atm, using a 2-fs integration step. Coordinates were recorded every 20 ps and then used for analysis, which in most cases involved Python scripts based on MDAnalysis [57]; lipid orientation vectors **n**_*i*_ were defined as in earlier work [26, 36]; lipid interdigitation was computed using MOSAICS [58]. Simulation snapshots were rendered with VMD [59].

### Potentials-of-mean-force (PMF) of membrane bending

PMF profiles as a function of applied membrane curvature were calculated as described elsewhere [25]. Briefly, one-dimensional density profiles of the lipid phosphate atoms were used to construct three-dimensional density maps, one for each leaflet, in a state of zero curvature. Sinusoidal deformations given by cos(2*πx/L*_*x*_) were then applied to each leaflet’s map to produce additional maps *ϕ*_*k*_(**x**) that represent a series of curved bilayer states. The latter maps were then used to define the Multi-Map collective variable used to induce a deformation of the membrane [25]. That is:

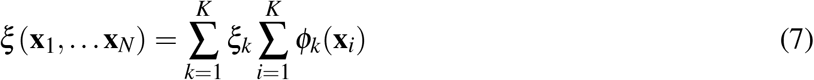

where **x**_*i*_ are coordinates of the phosphate atoms and *ξ*_*k*_ are constants quantifying the amount of curvature in each map. Note that *ξ* is directly proportional to 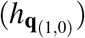 for a sinusoidal shape [60, 25]. More complex shapes are also supported by the Multi-Map method [25, 5], but a sinusoid allows for simpler analytical expressions of the relevant vector fields (Eq. 3). Cholesterol molecules were not included in the definition of *ξ*, as they might migrate between leaflets; in post-hoc analysis, we verified that the lateral distribution of cholesterol is not significantly correlated with curvature for any of the compositions used (Fig. 5).

To initialize the PMF calculations, values of *ξ* were derived from unbiased MD trajectories and collected into histograms with spacing *≈* 0.2 times the standard deviation of *ξ*. Random snapshots from each histogram bin were extracted from those trajectories and used to initialize umbrella-sampling windows [31]. Using uncorrelated initial snapshots improves statistical sampling compared to previous computations with Multi-Map [25]. All the umbrella-sampling windows were simulated concurrently using NAMD and the Colvars module [61], with harmonic restraints on *ξ* of force constant 0.6 kcal/mol for *≈* 1 *μ*s (Table 1). Potentials-of-mean-force (PMF) were extracted using the WHAM [62] and FCAM [63] methods, with no significant differences between the two methods. To maximize the computing time dedicated to examine the cholesterol-enriched bilayers, the umbrella-sampling calculations for the 1800-lipid POPC and DOPC pure-lipid membranes used shorter sampling times than for the other systems, which were nevertheless sufficient to observe convergence of their structural changes; the PMF curves in those two cases were however derived from unbiased histograms of *ξ*.

## Acknowledgments

This research was supported by the Divisions of Intramural Research of the National Institute of Neurological Disorders and Stroke (G.F. and L.R.F, NS003139) and the National Heart, Lung and Blood Institute (J.D.F.-G.). Computer resources were in part provided by NIH, through the “Biowulf” supercomputer, and in part by the National Center for Supercomputing Applications, through the “Blue Waters” supercomputer (funded by National Science Foundation awards OCI-0725070 and ACI-1238993, the State of Illinois, and the National Geospatial-Intelligence Agency). The authors thank Blue Waters team members James Phillips and William Kramer for their outstanding support.

