## Supporting Information for "Membrane free-energy landscapes derived from atomistic dynamics explain nonuniversal cholesterol-induced stiffening"

| | **MD, bending fluctuations (this work)** | | | **MD, tilt fluctuations [1]** | | | **X-ray diffraction [2]** | | |
|---|---|---|---|---|---|---|---|---|---|
| | POPC | | | | | | | | |
| $k_c$ (kcal/mol) | 17.0 | [13.5, | 22.1] | 19.1 | [17.9, | 20.3] | 11.9 | [9.3, | 14.5] |
| $k_t$ (kcal/mol/Å$^2$) | 0.102 | [0.084, | 0.127] | 0.079 | | | 0.099 | [0.051, | 0.147] |
| | POPC/CHOL | | | | | | | | |
| $k_c$ (kcal/mol) | 44.4 | [23.7, | 99.8] | ****$P$ = 0.002** | | | | | |
| $k_t$ (kcal/mol/Å$^2$) | 0.085 | [0.072, | 0.101] | $P$ = 0.16 | | | | | |
| | DOPC | | | | | | | | |
| $k_c$ (kcal/mol) | 18.0 | [13.0, | 26.2] | 17.1 | [15.9, | 18.3] | 11.6 | [10.8, | 12.4] |
| $k_t$ (kcal/mol/Å$^2$) | 0.103 | [0.080, | 0.138] | 0.092 | | | 0.128 | [0.116, | 0.140] |
| | DOPC/CHOL | | | | | | | | |
| $k_c$ (kcal/mol) | 28.5 | [16.0, | 60.3] | $P$ = 0.24 | | | | | |
| $k_t$ (kcal/mol/Å$^2$) | 0.081 | [0.064, | 0.106] | $P$ = 0.20 | | | | | |

**Table S1:** Bending moduli $k_c$ and tilt moduli $k_t$ estimated from membrane bending fluctuations. The estimated mean of each parameter, obtained by fitting Eq. 5 against the data, is reported along with the corresponding 95% confidence interval (CI) in brackets. Bending fluctuation results are from this work (Fig. 7), with uncertainty estimated by parametric bootstrapping [3]. The same distributions were used to perform statistical tests against the hypothesis that cholesterol has no effect, with the resulting $P$-values reported in the table. Also reported are previous results for the pure lipids using tilt fluctuations [1] and X-ray diffraction [2], with 95% CIs calculated from the reported standard errors, if available.

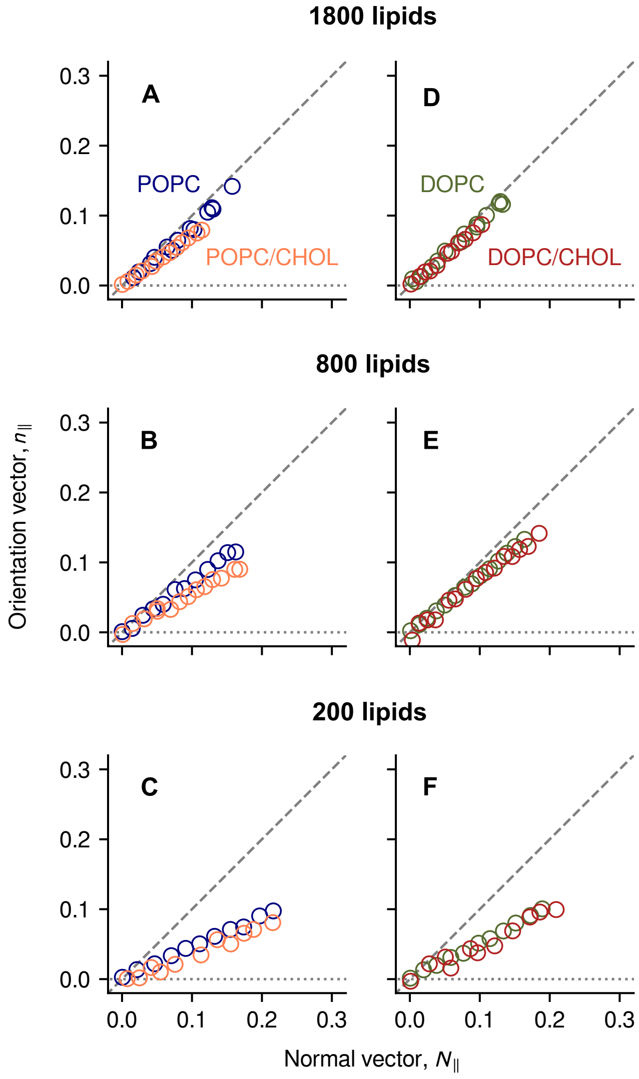


**Figure S1:** Change in lipid orientation as a function of curvature. The longitudinal component of the mean lipid orientation vector $n_{\|}$ was regressed onto the longitudinal component of the membrane normal vector, $N_{\|}$. Based on this regression, the value of $n_{\|}$ corresponding to the largest value of $N_{\|}$ in each bilayer is plotted for each umbrella-sampling simulation window. Grey dashed lines indicate the Helfrich-Canham theoretical prediction ($n_{\|} = N_{\|}$); dotted lines indicate no changes in lipid orientation ($n_{\|} = 0$). Results are shown for POPC and POPC/CHOL bilayers *(A-C)* and DOPC and DOPC/CHOL bilayers *(D-F)*. The values of the regression slopes are plotted in Fig. 3D,E and Fig. S2.

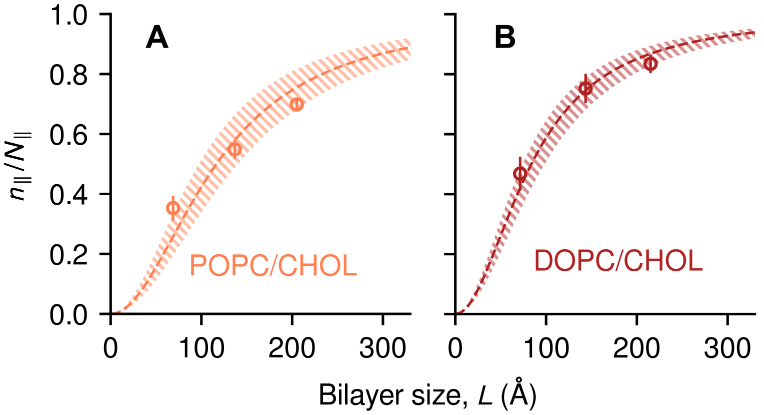


**Figure S2:** Analysis of the balance between bending and tilt energies for mixed bilayers. The expression in Eq. 3 is fitted against the values of $n_{\|}/N_{\|}$ for POPC/CHOL *(A)* and DOPC/CHOL bilayers *(B)*, using $k_c/k_t$ as the single fitting parameter. Dashed lines indicate best-fit curves, and striped bands the 95% CI around each curve.

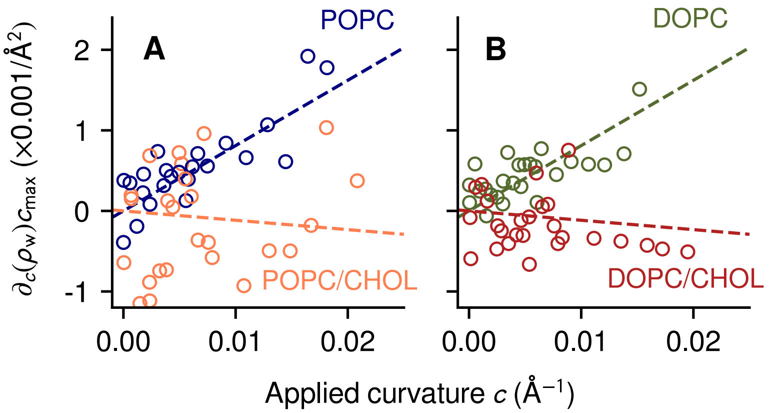


**Figure S3:** Changes in number of water molecules $\rho_w$ embedded in the hydrophobic section of each bilayer quantified for POPC and POPC/CHOL *(A)*, and DOPC and DOPC/CHOL *(B)*. Dashed lines indicate the results of linear fits of $\Delta\rho_w$ vs. the curvature $c$, performed on aggregated data from POPC and DOPC (slope = 0.081 ± 0.005 Å$^{-2}$) or from POPC/CHOL and DOPC/CHOL (slope = -0.011 ± 0.009 Å$^{-2}$). No significant differences were detected between POPC and DOPC or between POPC/CHOL and POPC/CHOL.

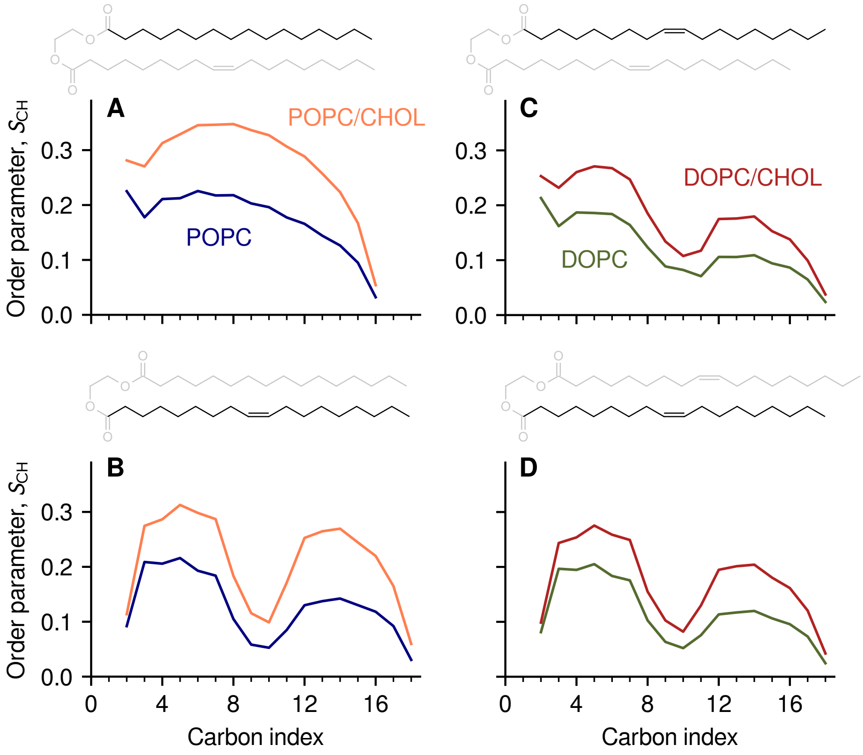


**Figure S4:** Atomic order parameters $S_{\mathrm{CH}}$ between acyl chain carbons and their bound hydrogens, computed as a function of the carbon atom's position along each chain. Results are shown for the *sn*-1 *(A)* and *sn*-2 chain *(B)* of POPC molecules, and for the *sn*-1 and *sn*-2 chains of DOPC molecules *(C-D)*. The chemical structure of each chain is shown alongside the order parameters' values with the same scale.

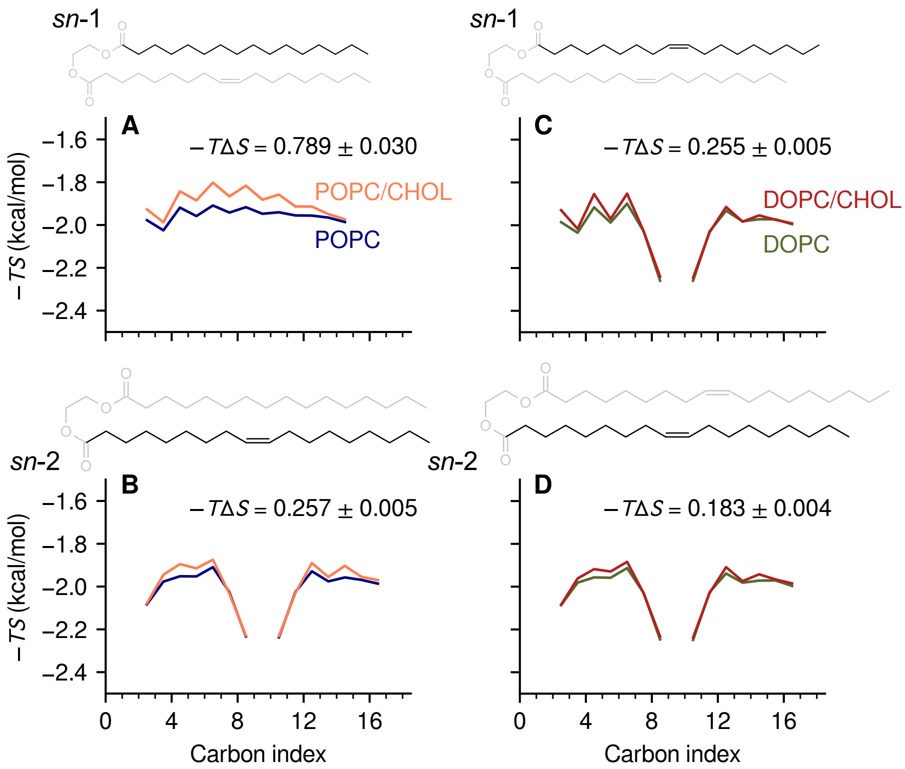


**Figure S5:** Entropy of lipid acyl chains, extracted from histograms of the torsional angles of carbon atoms with a spacing of 5°. The chemical structure of each acyl chain is shown alongside the reported entropy values as a function of the position along the chain of the second carbon atom of the torsional angle. Entropy values are shown as free-energy contributions, $-TS$, in kcal/mol units. Results are shown for the *sn*-1 *(A)* and *sn*-2 chain *(B)* of POPC molecules, for the *sn*-1 and *sn*-2 chains of DOPC molecules *(C-D)*. For each chain, the aggregated change $\langle -T\Delta S \rangle$ upon adding cholesterol is also reported in the panel (in kcal/mol units).

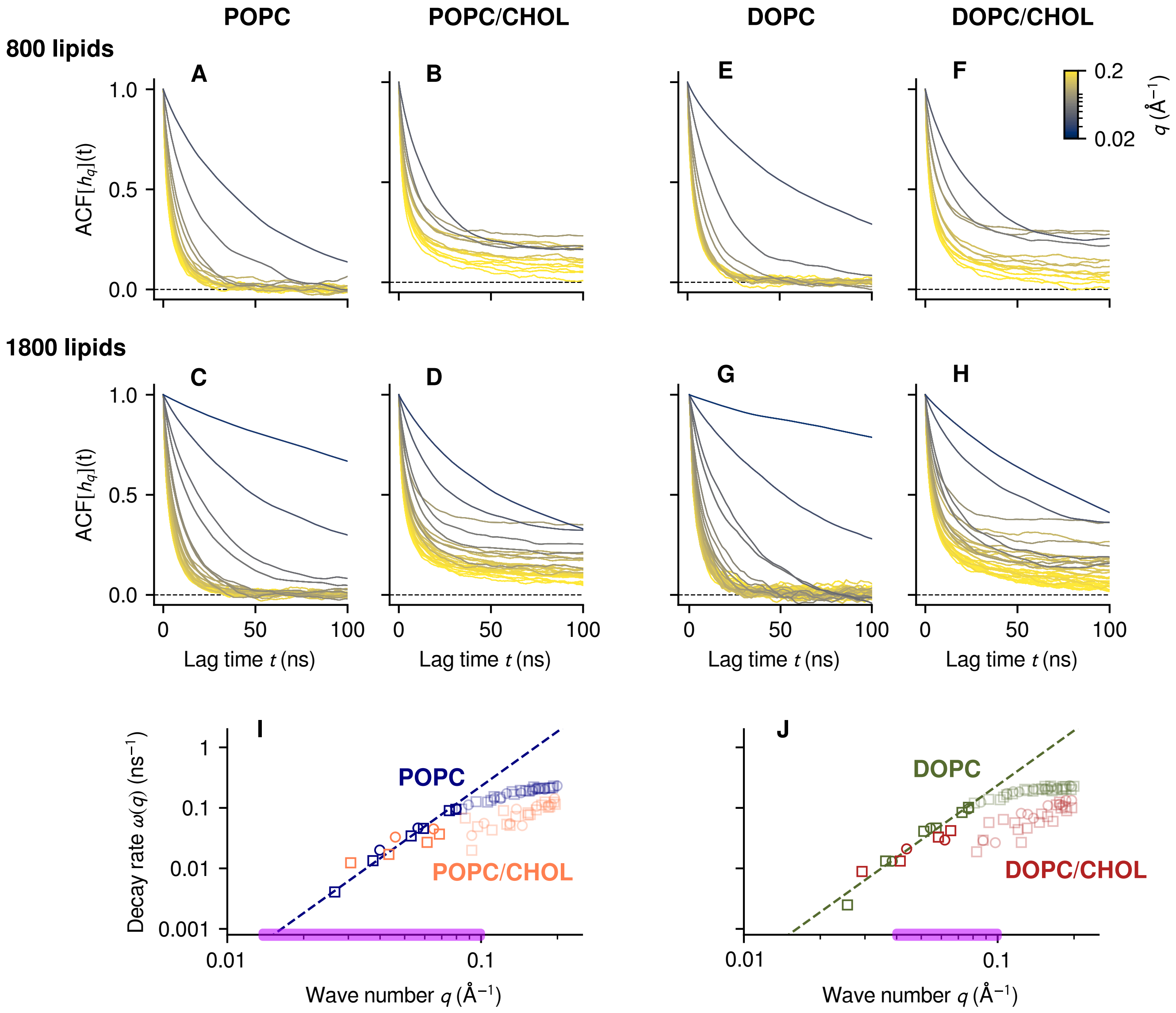


**Figure S6:** *(A-H)* Auto-correlation functions (ACFs) of the bending Fourier coefficients $h_\mathbf{q}$ for multiple bilayers, colored according to the value of $q$ as in the scale shown. *(I,J)* Decay rates $\omega(q)$, obtained by fitting ACFs to single exponentials, for POPC (blue), POPC/CHOL (orange), DOPC (green) and DOPC/CHOL (red). Decay rates from 800-lipid and 1800-lipid bilayers are shown as circles and squares, respectively; values for $q > 0.08$ Å$^{-1}$ are shown in transparent colors. Dashed lines indicate a linear fit for DOPC and POPC of $\omega(q)$ vs. $q^3$ as predicted by dynamic theories [4, 5, 6, 7] based on the Helfrich-Canham model [8, 9]. The intervals of $q$ where NSE experiments were carried out [10, 11] are highlighted in purple.

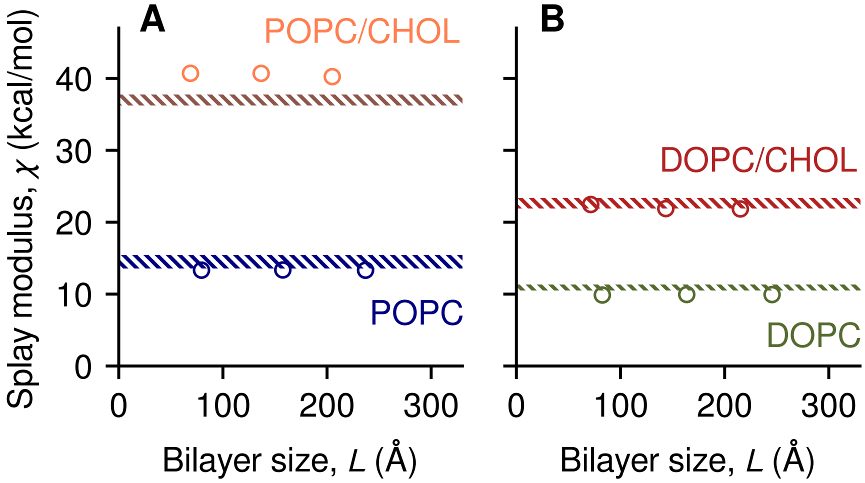


**Figure S7:** Lipid splay modulus $\chi$ computed from fluctuations of the mutual angles $\mathbf{n}_i \cdot \mathbf{n}_j$ between orientation vectors of lipid molecules [12]. *(A)* Values of $\chi$ for POPC (blue) and POPC/CHOL (orange) bilayers shown as circles; dashed bands areas indicate the 95% CIs of literature values [12]; because no values for POPC/CHOL were reported, data for the similar mixture POPC/POPS/CHOL 34:30:36 [12] are shown in brown. *(B)* Values of $\chi$ for DOPC and DOPC/CHOL, compared to published 95% CIs [11].
